# Increased expression of mitochondrial dysfunction stimulon genes affects chloroplast redox status and photosynthetic electron transfer in Arabidopsis

**DOI:** 10.1101/696740

**Authors:** Alexey Shapiguzov, Lauri Nikkanen, Duncan Fitzpatrick, Julia P. Vainonen, Arjun Tiwari, Richard Gossens, Saleh Alseekh, Fayezeh Aarabi, Olga Blokhina, Klará Panzarová, Zuzana Benedikty, Esa Tyystjärvi, Alisdair R. Fernie, Martin Trtílek, Eva-Mari Aro, Eevi Rintamäki, Jaakko Kangasjärvi

**Affiliations:** Organismal and Evolutionary Biology Research Programme, Faculty of Biological and Environmental Sciences, University of Helsinki, FI-00014 Helsinki, Finland; Viikki Plant Science Center, University of Helsinki, FI-00014 Helsinki, Finland; Institute of Plant Physiology, Russian Academy of Sciences, 127276 Moscow, Russia; Department of Biochemistry / Molecular Plant Biology, University of Turku, FI-20014 Turku, Finland; Turku Bioscience Centre, University of Turku, FI-20014 Turku, Finland; Max-Planck Institute for Molecular Plant Physiology, D-14476 Potsdam-Golm, Germany; Center of Plant Systems Biology and Biotechnology, 4000 Plovdiv, Bulgaria; Photon Systems Instruments, 664 24 Drásov, Czech Republic

**Keywords:** Arabidopsis thaliana, mitochondrial dysfunction stimulon, alternative oxidases, hypoxia, reactive oxygen species, photosynthetic electron transfer

## Abstract

Mitochondrial retrograde signals control expression of nuclear mitochondrial dysfunction stimulon (MDS) genes. Although MDS gene products mostly affect mitochondrial functions, they also influence production of reactive oxygen species (ROS) and redox status of chloroplasts. To study this inter-organellar interaction, we analysed the response of the Arabidopsis MDS-overexpressor mutant *rcd1* to methyl viologen (MV), which catalyses electron transfer from Photosystem I (PSI) to molecular oxygen, generating ROS in Mehler’s reaction. The response of plants to MV was investigated by imaging chlorophyll fluorescence in aerobic and hypoxic environments, and by membrane inlet mass spectrometry. Hypoxic treatment abolished the effect of MV on photosynthetic electron transfer in *rcd1*, but not in wild type. A similar reaction to hypoxia was observed in other MDS-activating lines and treatments. This suggests that MDS gene products contribute to oxygen depletion at the PSI electron-acceptor side. In unstressed growth conditions this MDS-related effect is likely masked by endogenous oxygen evolution and gas exchange with the atmosphere. In *rcd1*, altered Mehler’s reaction coincided with more reduced state of the chloroplast NADPH-thioredoxin oxidoreductase C (NTRC) and its targets, suggesting that NTRC performs feedback control of photosynthesis. This regulation may represent a novel mechanism whereby mitochondrial retrograde signalling affects chloroplast functions.

## Background

Photosynthetic light reactions are subject to precise transcriptional and post-translational regulation. Transcriptional regulation is, at least in part, mediated by retrograde signals that are emitted by the organelles to trigger nuclear transcriptional reprogramming. Post-translational regulation allows rapid operational control of photosynthesis in a changing light environment. For example, light-dependent acidification of the thylakoid lumen activates protective adaptation in two photosynthetic complexes, Photosystem II (PSII) and b_6_f. Protonation of the PSII subunit PsbS induces non-photochemical quenching (NPQ) [1]. NPQ allows PSII to convert light energy to heat rather than charge separation, thus protecting downstream components of photosynthetic electron transfer (PET) from excessive reducing power. In addition, acidification of the thylakoid lumen hinders electron transfer through the b_6_f complex in a process referred to as “photosynthetic control” [2, 3]. This also protects the downstream PET components, mainly Photosystem I (PSI), from excessive reducing power. When cellular energy demands increase, the activity of the thylakoid ATP synthase counterbalances acidification of the thylakoid lumen, thus lowering NPQ and easing the b_6_f control [4, 5].

These and many other adaptations of photosynthesis partly depend on thiol redox regulation. Numerous chloroplastic thiol regulatory enzymes receive reducing power either from ferredoxin through ferredoxin:thioredoxin disulfide oxidoreductase (FTR) or from NADPH through NADPH-thioredoxin oxidoreductase of type C (NTRC) [6–8] and relay this reducing power to thiol redox enzymes. This allows adjustment of metabolic processes in chloroplasts in accordance with light conditions and the redox status of the PET chain. Recently, several studies have revealed a functional link between redox states of chloroplast thiol enzymes and reactive oxygen species (ROS) [9–12]. Chloroplastic ROS were suggested to oxidise the abundant thiol enzymes 2-Cys peroxiredoxins (2-CPs), thus draining reducing power away from the thiol redox system.

One of the main sources of chloroplastic ROS is the reduction of molecular oxygen by PSI, referred to as Mehler’s reaction [13, 14]. This electron transfer pathway can be artificially enhanced by treating plants with the herbicide methyl viologen (MV, also known as paraquat), a compound that shuttles electrons from PSI to O_2_. The effect of MV on photosynthesis is observed at several levels. Firstly, MV contributes to oxidation of PSI, thus affecting PET [15, 16]. Secondly, the chemical allows Mehler’s reaction to out-compete other electron transfer pathways downstream from PSI, including cyclic electron flow [17], the pathways to FTR and NADPH, and further to thiol redox enzymes [18]. These two effects take place as soon as plants pre-treated with MV are exposed to light. Finally, MV stimulates the formation of ROS. Gradual light-dependent increase in ROS production rate ultimately leads to destabilization of PSII and to cell death [9, 18, 19].

Chloroplast ROS production and processing are sensitive to the expression of proteins encoded by the nuclear mitochondrial dysfunction stimulon (MDS) genes [9, 20, 21]. Expression of these genes is controlled by the retrograde signal triggered by perturbations of mitochondrial electron transfer. In Arabidopsis, the MDS signalling pathway is mediated by at least two transcription factors, ANAC013 [20] and ANAC017 [22], and is inhibited by the nuclear co-regulator protein RCD1 [9]. As expected, most proteins encoded by MDS genes are related to mitochondrial functions. In plants with enhanced MDS gene expression, including the *rcd1* mutant and *ANAC013* overexpressor, changes in mitochondria coincide with increased tolerance to MV activity in the chloroplasts [9, 20, 21]. Furthermore, chloroplasts of the *rcd1* mutant have altered redox state when compared to the Col-0 wild type [23, 24]. However, the molecular nature of this inter-organellar interaction remains obscure. One prominent set of MDS gene products are alternative oxidases (AOXs), mitochondrial enzymes with ubiquinol:oxygen oxidoreductase activity. These enzymes are likely candidates for a role in modulating chloroplastic ROS processing. AOXs are known to provide an extra-chloroplastic electron sink for PET [25–28]. Pharmacological or genetic inhibition of AOX activity correlated with suppressed photosynthesis and decreased tolerance of plants to MV [9, 27, 29]. Importantly, AOX activity has also been implicated in the maintenance of mitochondrial oxygen homeostasis [30, 31] and generation of hypoxia in developing plant tissues [32]. It has been suggested that AOXs have evolved as oxygen-scavenging enzymes that affect energy metabolism and mitochondrial ROS formation though decreasing intracellular concentrations of molecular oxygen [30, 33].

Studies using the Arabidopsis *rcd1* mutant suggested that AOX activity modulates PET [9], however the mechanistic details are yet unknown. Here we aimed to understand how enhancement of Mehler’s reaction by MV affects PET, and by which mechanisms mitochondrial retrograde signals regulating AOXs could influence these chloroplastic processes. Our data suggest that activated MDS signalling may contribute to depletion of molecular oxygen, thereby possibly affecting Mehler’s reaction. This putative pathway would represent a novel mode of interaction between mitochondria and chloroplasts.

## Results and discussion

### Methyl viologen inhibits photosynthetic oxygen evolution through fast generation of NPQ

Methyl viologen (MV) catalyses the transfer of electrons from Photosystem I (PSI) to molecular oxygen, resulting in oxidation of photosynthetic electron transfer (PET) chain and generation of reactive oxygen species (ROS). Increased ROS production gradually inhibits Photosystem II (PSII). The impact of catalytic concentrations of MV on PET is evident within the first seconds of illumination. To visualize this effect, we pre-treated wild-type (Col-0) Arabidopsis leaf discs with 1 μM MV overnight in darkness and imaged fast chlorophyll fluorescence rise induced by saturating light (OJIP, standing for F<u>o</u>, F<u>j</u>, F<u>i</u>, and F<u>p</u> = Fm phases of fluorescence induction kinetics [34, 35]). We performed the measurement without prior light exposure to minimize ROS-induced damage of PSII. The effect of MV was visible already after 40 msec of illumination as lowered Fi-Fm phase of the OJIP kinetics (Fig. 1A). This phenomenon has previously been ascribed to oxidative action of MV on the electron-acceptor side of PSI [15–17]. It has been proposed that MV releases “a traffic jam of electrons caused by a transient block at the acceptor side of PSI” [15]. Accordingly, the quantum yield of the electron transport flux until the PSI electron acceptors, defined as φRE1o = 1 − Fi/Fm [35], was diminished by MV treatment in all the tested lines (Fig. 1B).

**Figure 1.**
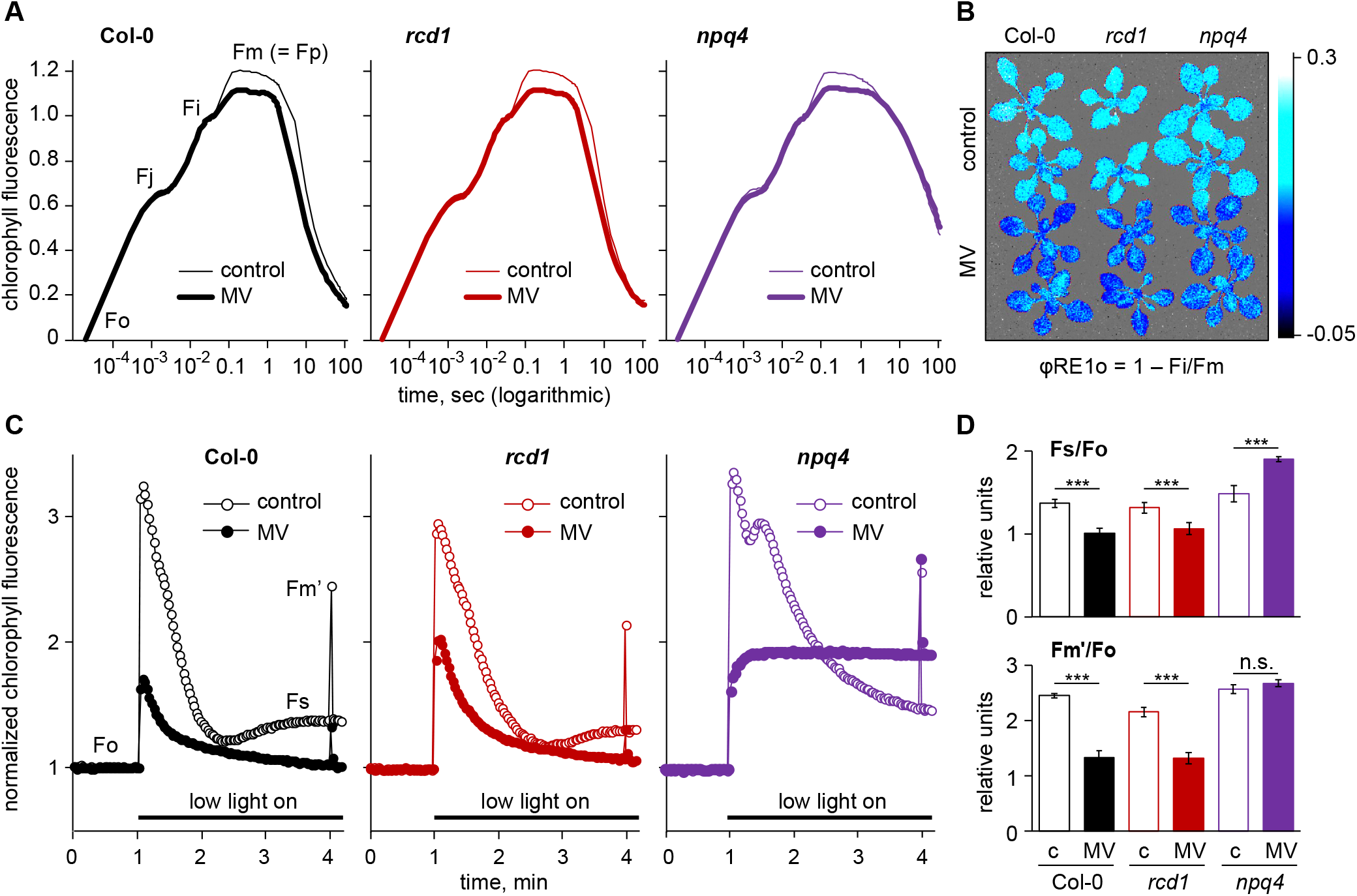
MV stimulates NPQ in the first minutes of illumination. (A) Kinetics of chlorophyll fluorescence excited by saturating light in leaf discs that were pre-treated with 1 μM MV in darkness. When plotted against time in logarithmic scale, the kinetics reveals several inflections defined as Fo-Fj, Fj-Fi, and Fi-Fm phases [34, 35]. Each phase is affected by different stages of PET. Fo-Fj relates to electron transfer within PSII up to Q_A_, Fj-Fi to intersystem redox states of PET, and Fi-Fm to electron transfer downstream from PSI [15, 16]. MV treatment lowered the Fi-Fm rise, indicating enhanced oxidation of PSI. Kinetics are double normalized to fluorescence at Fo and Fi (20 μsec and 40 msec, accordingly). The experiment was repeated three times with similar results. (B) False-colour image of the quantum yield of the electron transport flux until the PSI electron acceptors (φRE1o = 1 − Fi/Fm) in absence or presence of MV. (C) Kinetics of chlorophyll fluorescence during exposure to low light. Pre-treatment with MV decreased Fs and Fm’ in the wild type and in the MV-tolerant *rcd1*, but not in *npq4*. This suggests that fast effect of MV on PET is related to NPQ. All reads are normalized to Fo. (D) Quantification of chlorophyll fluorescence parameters described in (C). Untreated controls are labelled with “c”. Source data and statistical analyses are presented in Supplementary Table 1.

We next followed the dynamics of chlorophyll fluorescence over the first minutes of illumination with non-saturating light of 80 μmol m^−2^ s^−1^ (Fig. 1C). Wild-type leaf discs pre-treated with 1 μM MV and exposed to low light developed decreased steady-state fluorescence (Fs) and decreased maximal fluorescence under light (Fm’) as compared to the untreated control (Fig. 1C). The same was observed in the MV-tolerant mutant *rcd1*. In contrast, pre-treatment with MV did not change Fm’ in the *npq4* mutant and led to increase in Fs (Fig. 1C, D). The *npq4* mutant is deficient in the PSII subunit PsbS and is thus incapable of non-photochemical quenching (NPQ). Thus, the results indicated that in non-saturating light the quenching effect of MV on chlorophyll fluorescence in Col-0 and *rcd1* was related to NPQ. The increase of Fs in MV-treated *npq4* could possibly be due to the negative effect of thylakoid acidification on the electron transfer through the b_6_f complex, known as “photosynthetic control”.

Rapid development of NPQ / b_6_f complex control in MV-treated plants likely indicates enhanced acidification of thylakoid lumen under non-saturating light conditions. This acidification could be triggered by increased activity of the b_6_f complex and/or by suppressed proton efflux through the thylakoid ATP synthase [5]. The elevated b_6_f complex activity could result from increased electron flux downstream from PSI, while the ATP synthase could be suppressed due to competition of the enhanced Mehler’s reaction with thiol redox pathways that activate ATP synthase. However, validating these assumptions will be the subject of further research.

NPQ prevents charge separation in PSII. Therefore, it was reasonable to assume that MV would also inhibit O_2_ production by the PSII water-splitting complex. To test if this was indeed the case, we measured O_2_ evolution in MV-treated leaf discs using membrane inlet mass spectrometry (MIMS). This technique allows real-time monitoring of multiple compounds produced and absorbed through leaf gas exchange [36]. The use of isotope-labelled gases allowed us to distinguish production and absorption of gases (Suppl. Fig. 1). As expected, the measurements revealed markedly decreased O_2_ evolution already during the first minutes of illumination in the Col-0 wild type (Fig. 2A). Of note is the fact that MV did not inhibit respiration, as inferred from the measurements of CO_2_ evolution (Fig. 2A, Suppl. Fig. 1). The results suggest that upon exposure to light, MV-pre-treated wild type plants experienced fast activation of NPQ / b_6_f complex control, which led to suppression of PSII activity and decreased photosynthetic O_2_ evolution. This suppression of PSII activity occurred before any visible ROS-related damage to PSII.

**Figure 2.**
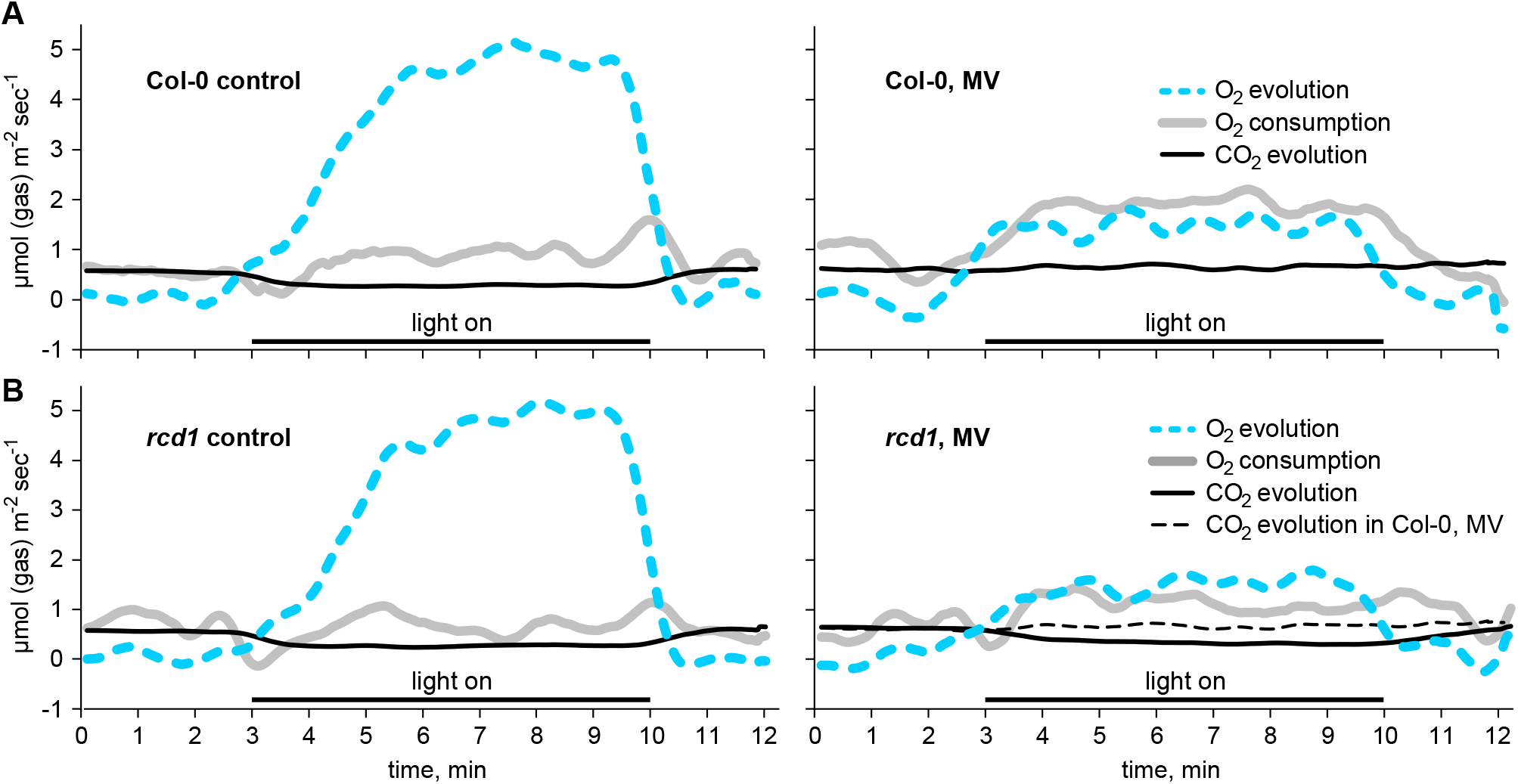
MV inhibits oxygen evolution in wild type and in *rcd1* in the first minutes of illumination. (A) MIMS measurements of O_2_ and CO_2_ gas exchange in the untreated (left) and MV pre-treated (right) Col-0 leaf discs. (B) O_2_ and CO_2_ gas exchange in the untreated (left) and MV pre-treated (right) leaf discs of *rcd1*. MV inhibited oxygen evolution both in Col-0 and *rcd1*, and CO_2_ reabsorption in Col-0, while respiration was unaffected in both lines. The full data set is presented in the Suppl. Fig. 1.

### Exposure to light reversibly inactivates MV in the *rcd1* mutant

We next analysed the effects of MV in the Arabidopsis mutant *rcd1*. In this mutant, constitutively activated mitochondrial retrograde signalling affects chloroplast ROS processing, resulting in tolerance to MV [9]. Similar to wild type, MV caused a pronounced decrease in O_2_ evolution in *rcd1*, which could be due to suppression of PSII electron-transfer activity by NPQ (Fig. 2B). Moreover, again as in Col-0, MV did not affect CO_2_ evolution in *rcd1*, suggesting uninhibited respiration. When plants are exposed to light, photosynthetic carbon fixation reabsorbs a fraction of CO_2_ produced by respiration, thus lowering net CO_2_ emission [37]. In control conditions, we observed this light-dependent change in both genotypes (Suppl. Fig. 1). Treatment with MV prevented light-dependent CO_2_ reabsorption in Col-0, but not in *rcd1* (Fig. 2B, right panel; Suppl. Fig. 1). This suggested that, in spite of NPQ-supressed PSII activity and oxygen evolution, photosynthetic carbon fixation was still active in MV-treated *rcd1*. This was possibly due to residual PSII activity and altered cyclic electron transfer pathways.

To address PET in *rcd1*, we measured the kinetics of chlorophyll fluorescence in low light. During the first minutes of low light exposure, *rcd1* performed like the wild type (Fig. 1C, D). However, longer light treatment led to gradual recovery of Fm’ in *rcd1*, but not in Col-0 (Fig. 3A). To test whether this effect was related to the release of NPQ, we next treated *rcd1* leaf discs with nigericin. This chemical impedes NPQ by preventing build-up of thylakoid proton gradient. When applied together with MV, nigericin suppressed both the initial drop of Fm’ and its concomitant recovery in *rcd1* (Suppl. Fig. 2A). We next generated an *rcd1 npq4* double mutant. When exposed to light, the MV pre-treated *rcd1 npq4* demonstrated neither the initial drop, nor the subsequent recovery of Fm’ characteristic of *rcd1* (Suppl. Fig. 2B). Importantly, during prolonged light treatment the tolerance of *rcd1 npq4* to MV-dependent PSII inhibition was significantly suppressed as compared to *rcd1* (Fig. 3B). Taken together, these observations suggest that alterations in NPQ contribute to MV tolerance of *rcd1* and that exposure to light gradually modified MV-induced NPQ in *rcd1*.

**Figure 3.**
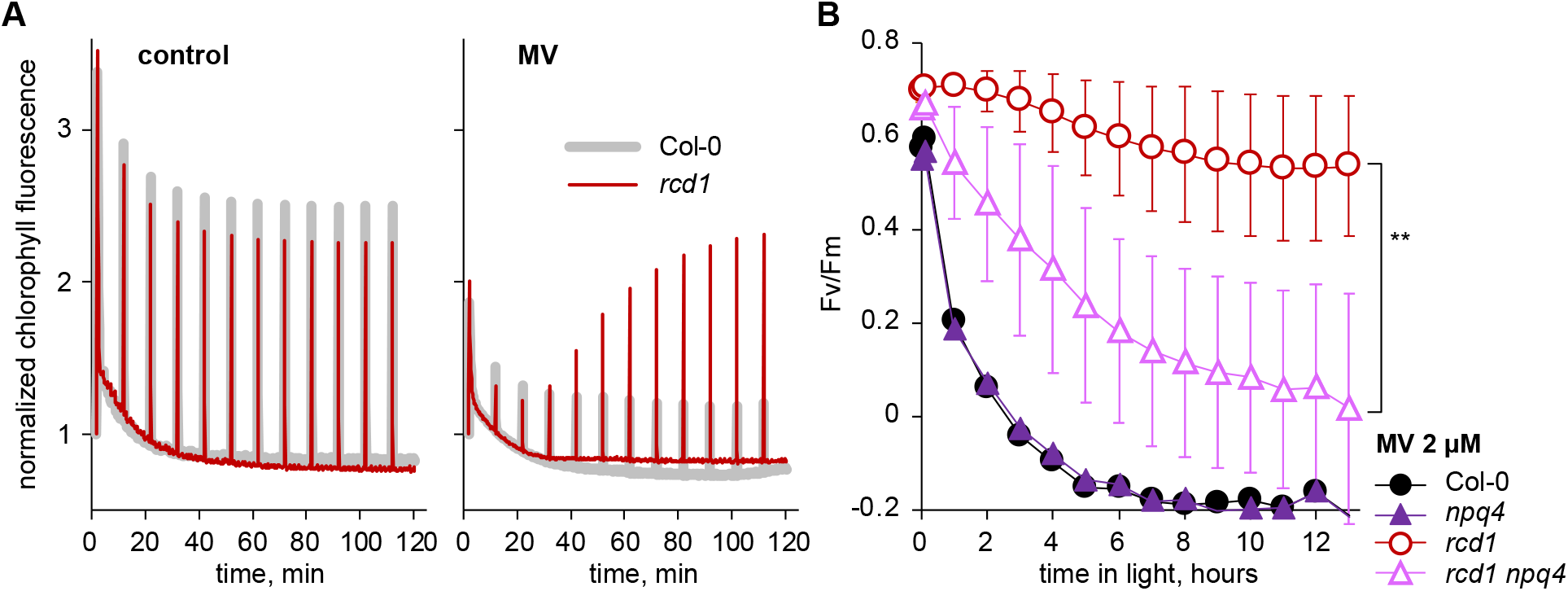
Exposure to light inhibits physiological activity of MV in the *rcd1* mutant. (A) Kinetics of chlorophyll fluorescence during 2 hours of exposure to low light. Saturating light pulses were triggered once in 10 minutes to measure Fm’. In *rcd1* pre-treated with MV (right panel), exposure to light resulted in gradual recovery of Fm’ to the control values (left panel), which was not observed in Col-0. The kinetics are normalized to Fo. The experiment was performed four times with similar results. The full experimental data set is presented in Fig. 7A. (B) The tolerance to MV-induced PSII inhibition was partially suppressed in the *rcd1 npq4* mutant as compared to *rcd1*, suggesting the importance of NPQ for MV tolerance of *rcd1*. Source data and statistical analyses are presented in Supplementary Table 1. The experiment was performed three times with similar results.

The tolerance of *rcd1* to MV is not due to diminished access of the chemical to PSI [9]. Thus, the described dynamics of chlorophyll fluorescence imply that light treatment gradually suppressed electron transfer through MV in the chloroplasts of *rcd1*, but not of Col-0 (Fig. 3A). Interestingly, this inactivation was reversible. This was shown by interrupting light treatment with 20-min dark periods. After each dark treatment, *rcd1* demonstrated decrease in Fm’ followed by recovery (Suppl. Fig. 2C), suggesting that the activity of chloroplastic MV was “reset” in darkness. This supports our assumption that in *rcd1* MV was not removed from its site of action, rather its function underwent reversible light-dependent quiescence.

### Physiological effect of MV in *rcd1* is abolished in hypoxic environment

The above data suggested that in the *rcd1* mutant exposure to light lowered the physiological activity of MV. The activity of MV depends on the availability of molecular oxygen. Treatment of plants with MV suppressed endogenous photosynthetic O_2_ evolution, but not respiration (Fig 2A, B). Moreover, formation of ROS, in particular H_2_O_2_, could form an oxygen sink further enhancing the oxygen deficit. This raises the question of whether these physiological circumstances would increase the demand for uptake of atmospheric oxygen. Thus, we exposed leaf discs to hypoxia by flushing nitrogen gas for 15 minutes in darkness, and measured chlorophyll fluorescence as in Fig. 1C. In all the tested lines hypoxia led to increased Fs and Fm’, as compared to the aerobic controls (Fig. 4A). This was anticipated, since molecular oxygen acts as an electron sink for a number of chloroplastic processes including the Mehler’s reaction and activity of the chloroplast terminal oxidase PTOX. In the wild type, MV markedly diminished the hypoxia-related rise in chlorophyll fluorescence (Fig. 4B). This was likely due to catalysis of Mehler’s reaction, which consequently compensated for oxygen deficiency. Importantly, the same effect of MV was observed in the *ptox* mutant, indicating that it was not associated with the PTOX activity (Suppl. Fig. 3). Similarly, MV lowered chlorophyll fluorescence in hypoxia-treated *npq4* and *stn7*, suggesting that the shift was not due to NPQ or chloroplast state transitions (Suppl. Fig. 3). In striking contrast to all of the above plant lines, in *rcd1* MV did not lower chlorophyll fluorescence under hypoxic conditions (Fig. 4A, B). This implied that hypoxia compromised the electron flow through MV in *rcd1*.

**Figure 4.**
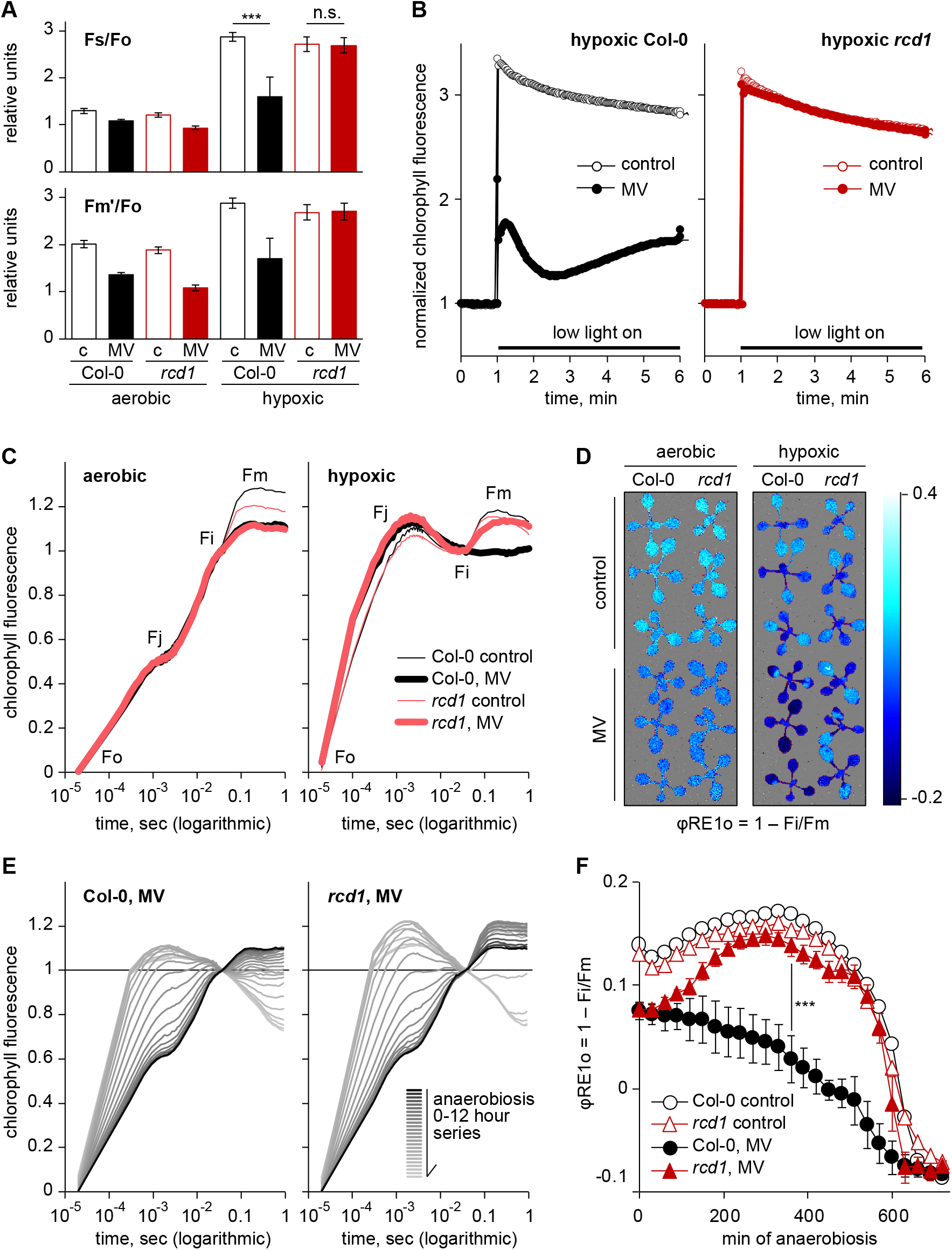
Hypoxic environment counteracts physiological effect of MV in the *rcd1* mutant. (A) Alterations in chlorophyll fluorescence induced by a 15-min pre-treatment with nitrogen gas in darkness. Source data and statistical analyses are presented in Supplementary Table 1. (B) Kinetics of chlorophyll fluorescence upon exposure to low light of leaf discs pre-treated with nitrogen as in (A). Under aerobic conditions, MV quenched fluorescence both in Col-0 and *rcd1*, while under hypoxia the effect of MV was not detectable in *rcd1*. The reads are normalized to Fo obtained in dark-adapted hypoxic conditions. Untreated controls are labelled with “c”. (C) Kinetics of chlorophyll fluorescence excited by saturating light. Leaf discs were pre-treated with 1 μM MV in darkness and imaged under aerobic (left panel) or hypoxic (right panel) conditions. The inhibitory effect of MV on the Fi-Fm phase was observed in both lines under aerobic conditions, but was absent from *rcd1* under hypoxia. Kinetics are double normalized to fluorescence at Fo and Fi (20 μsec and 40 msec, accordingly). (D) The same effect as in (C) shown with the false colour image of φRE1o = 1 − Fi/Fm. The experiment was performed four times with similar results. (E) Dynamic response of OJIP transients in MV-treated Col-0 and *rcd1* leaf discs subjected to hypoxia in AnaeroGen anaerobic gas generator. The experiment was performed twice with similar results. (F) The dynamics of φRE1o during transition to hypoxia described in panel (E). Source data and statistical analyses are presented in Supplementary Table 1.

It remained unclear whether the oxygen limitation affected MV activity directly at the electron-acceptor side of PSI, or indirectly, for example, through changes in mitochondrial respiration. To address this question, we performed OJIP imaging in leaf discs pre-treated overnight with 1 μM MV (Fig. 4C). Under aerobic conditions, the fluorescence induction kinetics was similar in Col-0 and *rcd1* (left panel). In both lines, MV lowered the Fi-Fm rise after 40 msec of illumination. Flushing nitrogen gas over leaf discs led to increased Fo-Fj phase in both lines. This effect has previously been attributed to induced fermentative metabolism and over-reduction of the chloroplast plastoquinone pool [38]. Most importantly, while suppression of the Fi-Fm rise by MV was still detected in the wild type, it was absent in *rcd1* (Fig 4C, right panel, Fig. 4D).

Fast change in OJIP kinetics induced by nitrogen gas flush made it difficult to observe the transition of PET from the aerobic to the hypoxic state. Thus, we generated hypoxia using an alternative approach, by placing MV-pre-treated leaf discs in the AnaeroGen anaerobic gas generator. This system decreases oxygen concentration below 0.5% while producing 9-13% of CO_2_ [39]. To prevent CO_2_ accumulation, we supplemented the anaerobic container with a CO_2_ absorbent reagent. Over 12 hours of dark incubation, 1-sec saturating light pulses were triggered once in 30 min to image OJIP kinetics. The treatment resulted in similar changes of OJIP as those observed with nitrogen gas flush. The raw fluorescence curves are presented in Fig. 4E, and the calculated φRE1o parameter in Fig. 4F. In MV-pre-treated *rcd1*, hypoxic treatment restored the Fi-Fm rise to the levels observed in MV-untreated controls, while this did not happen in MV-pre-treated Col-0 (Fig. 4E, F). Thus, physiological effect of MV on oxidation of the electron-acceptor side of PSI was prevented by hypoxia in the *rcd1* mutant.

These results raise the question why the *rcd1* mutant is more responsive to externally generated hypoxia than the wild type. At least two possibilities exist. One relates to altered stomatal functions of the mutant. Indeed, *rcd1* has been shown to have slightly higher stomatal conductance than the wild type in light [40]. However, this difference is unlikely to play a decisive role in darkness, when stomata should be largely closed, and in the presence of MV that has also been shown to promote stomatal closure [41]. The effect of hypoxia on the activity of MV was sustained in *rcd1* during several hours of dark hypoxic treatment, while it was completely absent from Col-0 (Fig. 4F), which is hard to explain by moderate difference in stomatal conductance. Another possible reason for the sensitivity of *rcd1* to hypoxia may be related to the altered mitochondrial functions of the mutant.

In a recent study, we demonstrated that expression of MDS genes activated by mitochondrial retrograde signalling, and subsequent accumulation of MDS gene products affected mitochondrial respiration in *rcd1* [9]. One group of the MDS gene products, AOXs, have been proposed to limit oxygen concentrations in plant mitochondria [30, 31] and tissues [32]. Taking this into account, it could be possible that the enhanced AOX activity may provide increased oxygen sink in the leaf tissues of *rcd1*. In standard growth conditions the effect is masked by gas exchange through stomata and photosynthetic oxygen evolution. Treatment with MV inhibits O_2_ evolution (Fig. 2A, B) and stimulates stomatal closure [41]. It could thereby promote oxygen depletion in *rcd1* tissues, thus inhibiting Mehler’s reaction. The relevance of this effect for MV tolerance of *rcd1* and, more generally, for the interaction between the energy organelles, is a subject of further research.

### Increased expression of MDS genes is linked to hypoxic response

We aimed to find out whether the altered response to hypoxia observed in *rcd1* exists in other MDS-inducing lines or treatments. The mitochondrial respiration inhibitor antimycin A (AA) activates MDS retrograde signalling. Accordingly, in wild-type plants pre-treated with AA, hypoxia led to decreased MV activity (Suppl. Fig. 4A, B). Measurement of OJIP kinetics in plants overexpressing *ANAC013* [20, 21] revealed quiescence of MV by hypoxia similar to that in *rcd1* (Suppl. Fig. 4C). These observations demonstrated that oxygen availability affected the MV response not only in the *rcd1* mutant, but also under other perturbations activating MDS gene expression.

To explore possible similarities of MDS gene expression with hypoxic response, we performed meta-analysis of the corresponding publically available transcriptomic datasets, including the genes differentially expressed in the *rcd1* mutant in standard growth conditions, the genes affected by a 3-hour treatment with 50 μM AA, or by a 2-hour treatment with hypoxia. A statistically significant overlap was found between the genes whose expression was activated in *rcd1*, under AA, and under hypoxic treatment (Fig. 5). The 19 genes that were activated in all the perturbations included the hypoxia-responsive universal stress protein 1 (*HRU1*), the stress-responsive transcription factor *ZAT10*, transcription factor *WRKY25*, as well as the MDS genes *AOXs* and *SOT12*. The SOT12 sulfotransferase is a component of the 3’-phosphoadenosine 5’-phosphate (PAP) metabolic pathway. PAP mediates retrograde signalling by mitochondria and chloroplasts and is linked to the activity of RCD1 [9, 42]. Noteworthy, RCD1 protein itself was shown to interact with transcription factors implicated in mitochondrial (ANAC013/ANAC017 [9]) and chloroplast (Rap2.4a [23]) retrograde signalling, along with dozens of other transcription factors [43]. This suggests that the role of RCD1 as the hub merging retrograde and other signalling pathways in the nucleus could be indirectly modulated by cellular oxygen availability. The full gene list is presented in Supplementary Table 2. These results suggested that transcriptional reprogramming induced by hypoxia bears similarity to the changes in gene expression observed in the *rcd1* mutant or after AA treatment (Fig. 5, Supplementary Table 2).

**Figure 5.**
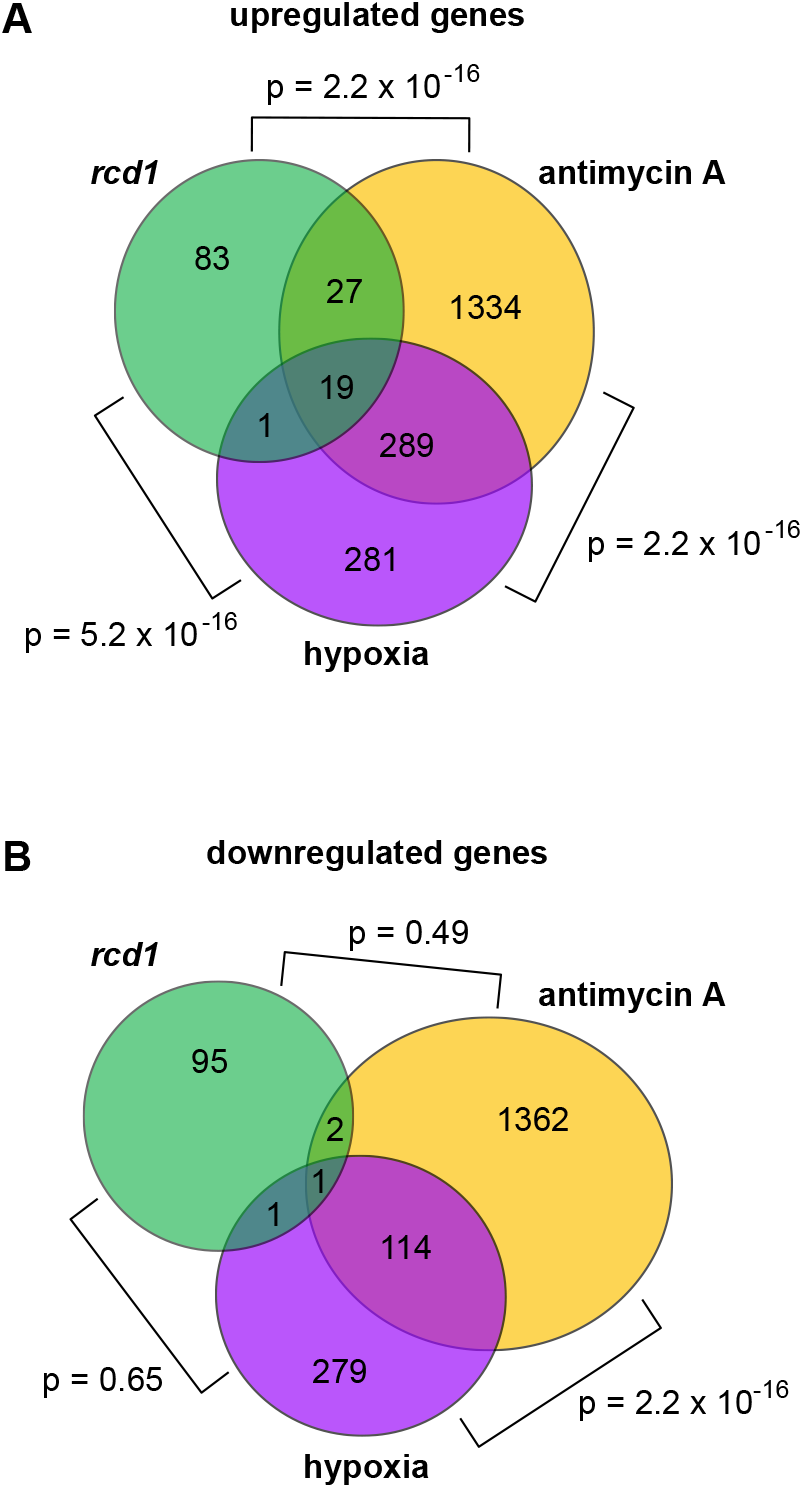
Similarities in transcriptional response of MDS-inducing perturbations and hypoxia. Analysis of publically available transcriptomic datasets obtained in the *rcd1* mutant, in wild-type plants treated with antimycin A (AA) or in wild types treated with hypoxia. Venn diagrams show the overlap of up- (A) and down-regulated (B) genes. Genes with at least 2-fold change in expression and a significance of p<0.05 were considered as up- or down-regulated. Statistical analysis was performed by a pairwise Fisher’s exact test.

Taken together, our results suggest that the activity of one or more MDS gene products may affect oxygen availability at the electron-acceptor side of PSI. The effect may likely be related to alterations in mitochondrial respiration and AOX activity. This opens up new experimental possibilities to explore the mechanisms and significance of chloroplastic and mitochondrial retrograde signalling in natural physiological conditions.

### Chloroplast NTRC system is over-reduced in *rcd1*

The above-described and previously published results indicated decreased ROS production in the chloroplasts of *rcd1* and other MDS-inducing mutants or treatments [9, 20, 21]. Chloroplastic ROS act as an electron sink for thiol redox enzymes [10–12]. Thus, suppressed ROS production likely results in more reduced redox state of these enzymes. Accordingly, chloroplast 2-Cys peroxiredoxin (2-CP) pool was more reduced in the *rcd1* mutant than in wild type [9]. The main enzyme reducing 2-CPs *in vivo* is NADPH-thioredoxin oxidoreductase of type C (NTRC) that is reduced by NADPH [6–8]. We assessed the *in vivo* redox state of NTRC in *rcd1* with thiol bond-specific labelling. Similarly to 2-CP, the NTRC pool was more reduced in *rcd1* than in wild type both in light and darkness (Fig 6A). This enzyme controls a number of chloroplastic processes including ROS processing [10, 11, 44], activities of thylakoid NADH dehydrogenase (NDH) complex mediating cyclic electron transfer [45], and of ATP synthase [46, 47].

**Figure 6.**
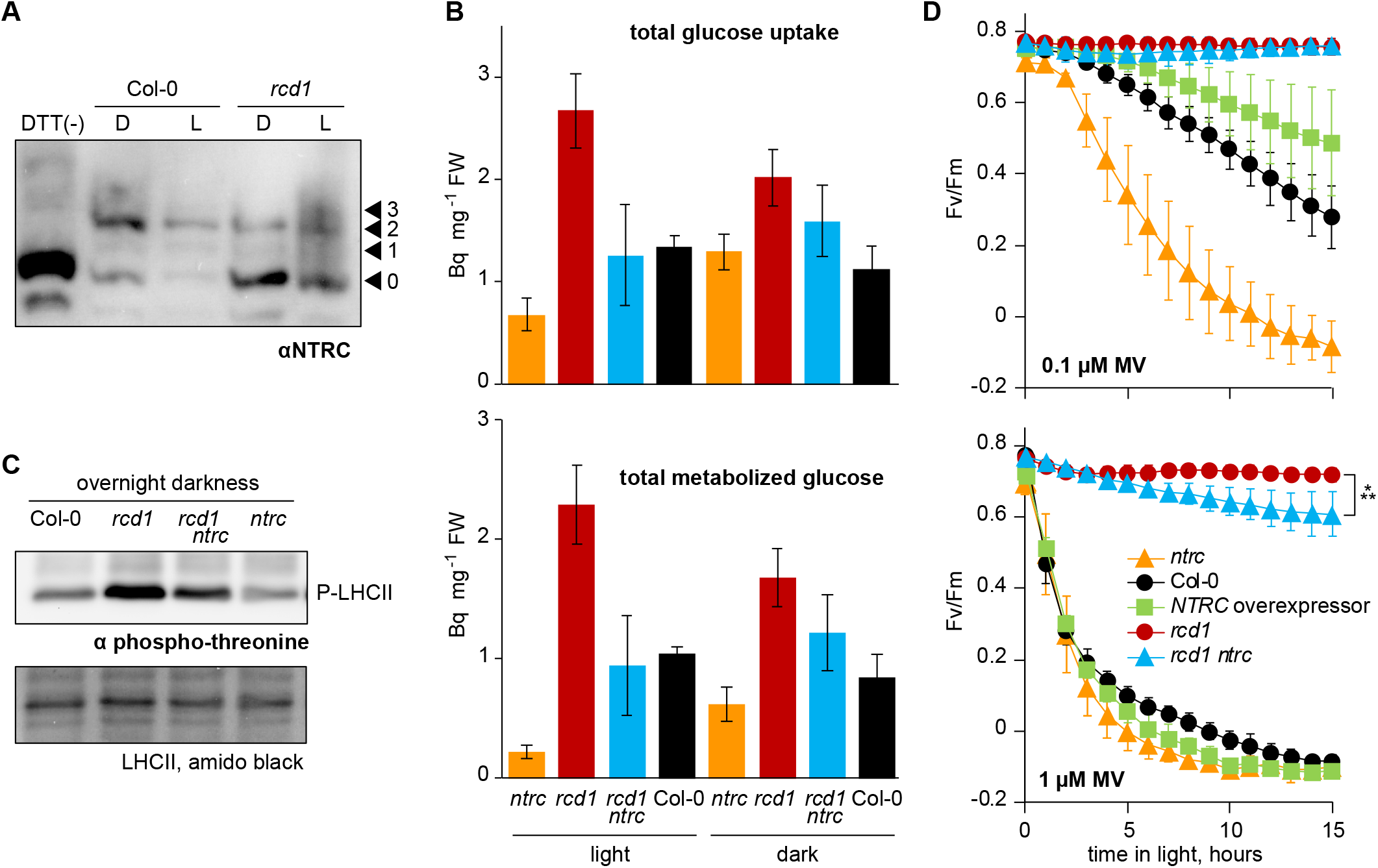
Chloroplast NTRC mediates MV response and other phenotypes of *rcd1*. (A) Chloroplast NTRC pool is more reduced in *rcd1* both in darkness (D) and light (L). Leaf protein extracts were subjected to thiol bond-specific labelling as described in [9]. Extracts were first treated with N-ethylmaleimide that blocked all the free thiol groups. Next, the *in vivo* thiol bridges were reduced with DTT. Finally, the extracts were treated with 5-kDa methoxypolyethylene glycol maleimide to label all the newly opened thiol groups. After separation in SDS-PAGE, the extracts were immunoblotted with the αNTRC antibody. The unlabelled form (0) corresponds to *in vivo* reduced, while the labelled forms (1, 2, 3) to *in vivo* oxidized fractions of NTRC. The experiment was performed twice with similar results. (B) Total metabolized and total consumed radiolabelled ^14^C glucose treated to light or dark-adapted rosettes. Mean values ± standard errors are presented. The full dataset is presented in the Supplementary Table 3. (C) Phosphorylation of LHCII in overnight dark-adapted seedlings as determined by immunoblotting with anti-phospho-threonine antibody. Amido black staining of total LHCII is shown in the lower panel as the loading control. (D) The tolerance to MV-induced PSII inhibition depends on NTRC. Two concentrations of MV were used, 0.1 μM MV (top panel), or 1 μM MV (bottom panel). The *ntrc* mutant was more sensitive and the *NTRC* overexpressor line more tolerant to MV as compared to Col-0. Tolerance to MV was partially suppressed in the *rcd1 ntrc* mutant as compared to *rcd1*. Source data and statistical analyses are presented in Supplementary Table 1. The experiment was performed three times with similar results.

To assess whether the phenotypes of *rcd1* depend on NTRC, we generated and characterized an *rcd1 ntrc* double mutant. To compare metabolic fluxes of *rcd1* and *rcd1 ntrc*, we fed leaf discs with uniformly ^14^C-labelled glucose and analysed the distribution of radioactive label between fractionated plant extracts and evolved CO_2_, as described in [9] (Supplementary Table 3). Total uptake and metabolism of glucose, significantly elevated in *rcd1* in light, were suppressed to wild-type levels in *rcd1 ntrc* (Fig. 6B). These results suggest that alterations in the energy metabolism of *rcd1* were partially mediated by NTRC.

NTRC regulates activity of the thylakoid NDH complex, one of the major pathways mediating chloroplast cyclic electron flow and reduction of plastoquinone pool in light and darkness. Hence, we next evaluated NDH activity in *rcd1* by assessing the redox state of the plastoquinone pool. Reduced plastoquinone pool activates the chloroplast kinase STN7 that phosphorylates the light-harvesting antenna complex II (LHCII). Thus, phosphorylation of LHCII can be used as an indirect measure of the plastoquinone redox state. Immunoblotting of total protein extracts from overnight dark-adapted seedlings with anti-phospho-threonine antibody revealed increased phosphorylation of LHCII in *rcd1* (Fig 6C). This indicated that similarly to *NTRC*-overepressing plants [45], the plastoquinone pool was reduced in *rcd1* in darkness, which is suggestive of increased NDH activity. Dark LHCII phosphorylation in *rcd1 ntrc* was suppressed as compared to *rcd1* (Fig. 6C). Thus, the increased activity of the NDH complex in *rcd1* was likely mediated by NTRC. Gas exchange measurements indirectly suggested altered cyclic electron flow in MV-treated *rcd1* (Fig. 2B). Further research is needed to address the roles played by NTRC and NDH in this response.

### NTRC is linked to chloroplast ROS processing and regulation of PET

Long illumination of MV-treated plants revealed that the *ntrc* mutant was more sensitive to MV than Col-0, and *rcd1 ntrc* was more sensitive to MV than *rcd1*. The *NTRC* overexpressor line [48] was more tolerant to MV than the wild type (Fig. 6D). We thus tested how NTRC was implicated in specific reactions of *rcd1* to MV described above. Leaf discs were pre-treated with MV in darkness and exposed to low light as in Fig. 3A (Fig. 7A). Treatment with MV resulted in NPQ-related decrease of Fm’ in all the tested lines. However, no *rcd1*-specific Fm’ recovery was observed in *rcd1 ntrc*. This indicates that the gradual release of NPQ in *rcd1* was dependent on NTRC. Acidification of the thylakoid lumen, and thus NPQ, is reversely correlated with the activity of thylakoid ATP synthase [5]. NTRC activates ATP synthase by thiol reduction [46]. Hence, it is likely that MV-induced thylakoid acidification was counteracted in *rcd1* through NTRC-dependent activation of ATP synthase. Of note, no changes in ATP synthase activity were detected in *rcd1* under standard light- or dark-adapted conditions (details in Suppl. Fig. 5). Further research is, however, needed to address the dynamics of ATP synthase activity in relation to the intensity of Mehler’s reaction.

**Figure 7.**
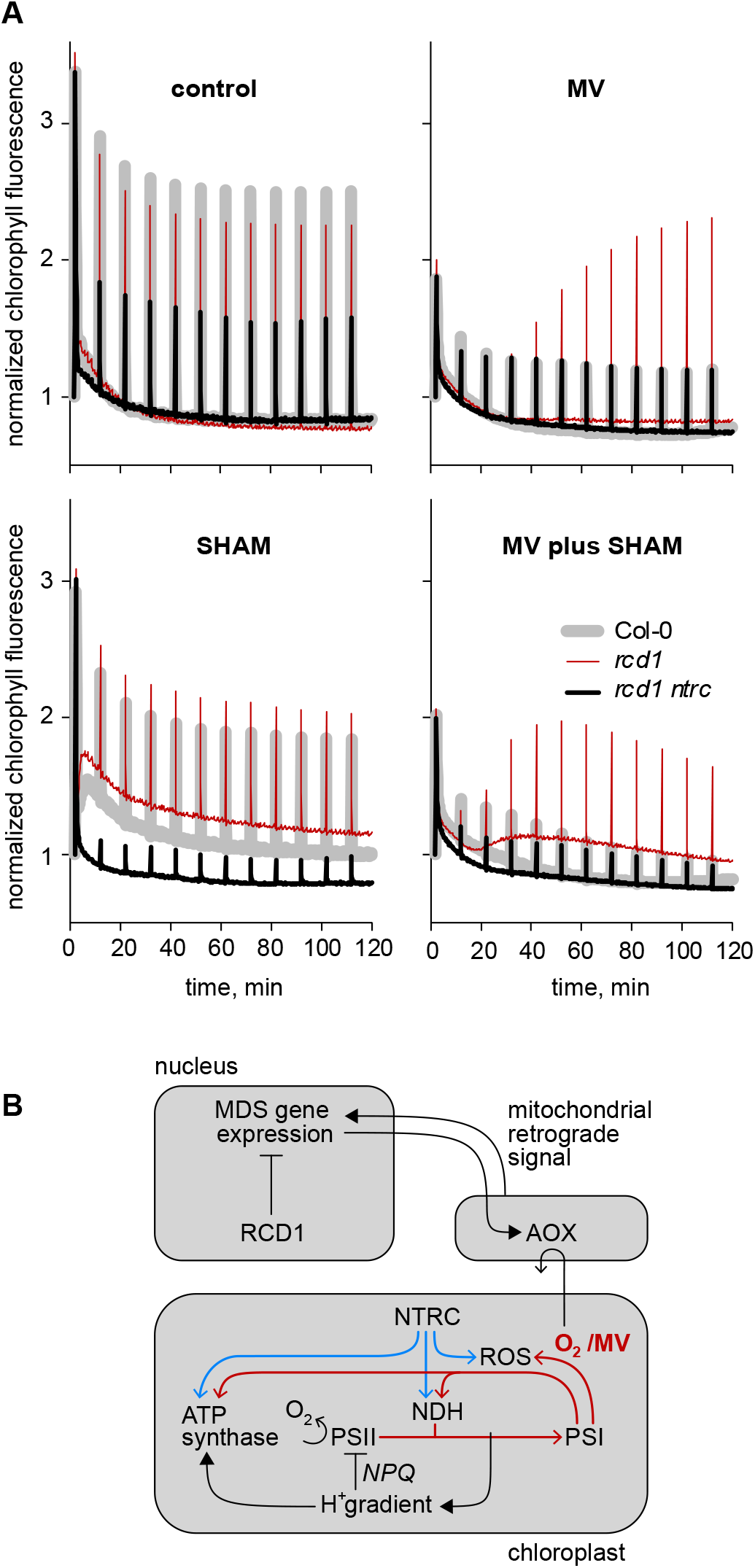
Interaction between mitochondria and chloroplasts in the MDS-inducing conditions. (A) Kinetics of chlorophyll fluorescence during 2 hours of exposure to low light. Saturating light pulses were triggered once in 10 minutes to measure Fm’. The *rcd1*-specific kinetics of Fm’ observed in MV-treated leaf discs was suppressed in *rcd1 ntrc*. The AOX inhibitor SHAM modified the response of PET to MV in *rcd1*, but not in *rcd1 ntrc*. The kinetics are normalized to Fo. The experiment was performed four times with similar results. (B) Altered mitochondrial respiration depends on the mitochondrial retrograde signal that activates expression of the nuclear MDS genes. The nuclear RCD1 protein suppresses expression of MDS genes. Thus, in the *rcd1* mutant MDS genes are constitutively induced. The activity of MDS gene products, including AOXs, may act as the sink for molecular oxygen. This can affect availability of O_2_ at the electron acceptor side of PSI, suppressing chloroplastic ROS production *via* Mehler’s reaction. Chloroplastic ROS act as the electron sink for the NTRC system. Through NTRC, altered chloroplastic ROS processing may affect PET. Relevant pathways of chloroplast electron transfer are shown with red arrows, NTRC-dependent redox control with blue arrows. By catalysing Mehler’s reaction, MV may drain reducing power away from ATP synthase. This promotes formation of proton gradient and onset of NPQ / b_6_f complex control inhibiting PSII electron transfer and O_2_ evolution activity. The resulting oxygen depletion reveals the putative effect of the MDS gene products on the chloroplastic Mehler’s reaction, contributing to MV tolerance of the *rcd1* mutant.

To test whether the activity of mitochondrial AOXs was implicated in this regulation, we pre-treated leaf discs with the AOX inhibitor salicylhydroxamic acid (SHAM). SHAM altered the response of *rcd1* to MV, resulting in increasing Fs after about 20 min of illumination and decreasing Fm’ after about 60 min (Fig. 7A). By contrast, no change in Fs was observed in *rcd1 ntrc*. Rising Fs may indicate increased reduction of the plastoquinone pool [9]. This observation hinted that inhibition of AOX activity caused an NTRC-dependent flow of reducing power to the plastoquinone pool. As NTRC controls the activity of thylakoid NDH complex [45], it is reasonable to suggest that the reduction of the plastoquinone pool was mediated by NDH. Hence, the mitochondrial AOX activity was likely linked to regulation of chloroplast thiol redox enzymes and thus PET. Overall, the above results indicated that activated expression of MDS genes coincided with a more reduced state of the chloroplast NTRC system and its targets, which affected the performance of PET at different levels.

The nuclear protein RCD1 likely plays multiple roles in the regulation of chloroplast thiol redox enzymes. Binding to ANAC013/ANAC017 transcription factors allows RCD1 to inhibit expression of MDS genes including *AOXs* [9]. Further control over organelles might be achieved through interaction of RCD1 with other transcription factors, such as Rap2.4a [23]. Finally, RCD1 protein is sensitive to ROS/redox-related retrograde signals emitted by the chloroplast [9]. Dissecting this trans-organellar regulatory loop is the subject of further research.

## Conclusions

Mitochondrial retrograde signalling controls expression of the nuclear MDS genes in Arabidopsis through the activity of at least two transcription factors, ANAC013 and ANAC017. The nuclear regulatory protein RCD1 interacts with these transcription factors, suppressing transcription of MDS genes [9]. Thus, MDS gene expression is constitutively increased in the *rcd1* knockout mutant. Accumulation of MDS gene products such as AOXs alter mitochondrial respiration. They also affect ROS metabolism and redox states of enzymes in the chloroplast (Fig. 7A) [9]. Here we show that the activity of one or more MDS gene products likely lowers oxygen availability in the chloroplasts (Fig. 7B). In standard growth conditions the effect is masked by photosynthetic oxygen evolution and gas uptake from the atmosphere. However, it can be observed in a hypoxic environment and following treatment with MV. The decreased oxygen availability at the electron-acceptor side of PSI coincides with, and possibly determines, the more reduced state of the chloroplast thiol enzyme NTRC and its targets, including the thylakoid ATP synthase and the thylakoid NDH complex. Through these components, oxygen limitation potentially modulates performance of PET (Fig. 7B). The proposed regulation may represent a novel mechanism, by which mitochondrial retrograde signalling affects photosynthesis.

## Materials and methods

### Plant lines and growth conditions

*Arabidopsis thaliana* (Col-0) plants were cultivated on soil (1:1 mix of peat and vermiculite) at a 12-hour photoperiod and light intensity of 220–250 μmol m^−2^ s^−1^. For the measurements of LHCII phosphorylation, seedlings were grown for 12 days on 1 x MS basal medium (Sigma-Aldrich) with 0.5% Phytagel (Sigma-Aldrich) without added sucrose, at a 12-hour photoperiod and light intensity of 150-180 μmol m^−2^ s^−1^. Arabidopsis *npq4-1* [49], *rcd1-4* (GK-229D11), *ptox* [50], *stn7* (SALK_073254), and *ntrc* (SALK 096776) mutants, as well as the *ANAC013* overexpressor line [20], and the *NTRC* overexpressor line [48], are of Col-0 background.

### Chemical and hypoxic treatments

For treatments with chemicals, cut leaf discs were placed on Milli-Q water with added 0.05% Tween 20 (Sigma-Aldrich), with or without MV. Unless specified otherwise, 1 μM MV was used. Final concentration of SHAM was 2 mM. It was prepared from the 2 M SHAM stock in DMSO. Thus, 1/1000 volume of DMSO was added to SHAM-minus controls. AA was applied in the final concentration of 2.5 μM, as discussed in [9]. Dark pre-treatment with MV lasted from 1 hour to overnight depending on the experiment. Pre-treatment with SHAM lasted for 1 hour. Pre-treatment with AA was overnight. To generate hypoxic environment, nitrogen gas was flushed inside a custom-built chamber that contained detached plant rosettes or a multi-well plate with floating leaf discs. Chlorophyll fluorescence imaging was performed through the top glass lid of the chamber. Alternatively, the multi-well plate with plant material was placed into the sealed plastic bag of the AnaeroGen Compact anaerobic gas generator system (Oxoid). In the same bag, resazurin Anaerobic Indicator (Oxoid) was placed to control for anaerobic conditions, and LoFloSorb non-caustic containing carbon dioxide absorbent (Intersurgical) to prevent accumulation of CO_2_.

### Direct fast imaging of OJIP chlorophyll fluorescence kinetics

The imaging of OJIP fluorescence kinetics was performed using FluorCam FC800F from Photon Systems Instruments, Czech Republic (www.psi.cz). The instrument is described in [51]. It contains the ultra-fast sensitive CMOS camera, TOMI 3, developed by Photon Systems Instruments, that performs image acquisition with maximum frame rate of 20 μsec. All acquired data are stored in the internal memory (1 GB) and transferred to the computer *via* the 1 GB communication Ethernet protocol. The required time resolution and minimization of the storage demands was achieved by the ability of the camera to record images on logarithmic or semi logarithmic time scale. FluorCam software was used to control the instrument and to analyze the data. The OJIP imaging protocol was comprised of the triple measurement of the background signal followed by three 20-μsec flashes of saturating light to measure Fo and then a 100-second (Fig. 1A) or a 1-second (all other figures) saturating light pulse (3 500 μmol m^−2^ s^−1^) to follow OJIP kinetics. Both the background and the Fo values were averaged for further calculations. The frame period was set at 300 μsec, and the integration time was 35-50 μsec. The excitation light was generated by a pair of blue LED panels (470 nm) and filtered by dichroic filters that block light at 490-800 nm to avoid crosstalk with the detection. To record chlorophyll fluorescence signal, the camera was equipped with 700-750-nm band filters.

### PAM chlorophyll fluorescence imaging

To measure kinetics of Fs and Fm’ and to assay long-term PSII inhibition, chlorophyll florescence was measured using Walz Imaging PAM essentially as described in [9].

### Measurement of gas exchange by membrane inlet mass spectrometry

For the gas exchange measurements, 14-mm leaf discs were floated overnight in Milli-Q H_2_O supplemented with Tween 20 with or without 1 μM MV at 20 °C. Following the overnight incubation, 12.5-mm discs were cut from the center of the pre-treated discs in very dim light, and loaded into a sealed MIMS cuvette calibrated to 22 °C. The cuvette was purged using air scrubbed of ^12^CO_2_ with carbosorb before ^13^CO_2_ gas (99% ^13^CO_2_, Sigma-Aldrich, USA) was injected to approximately 2% by volume, and ^18^O_2_ gas (98% ^18^O_2_, Cambridge Isotope Laboratories, UK) was enriched to approximately 3%. Samples were kept in darkness until gasses equilibrated between all areas of the leaf (approximately five minutes). Then data acquisition was commenced, comprising three minutes of darkness, seven minutes of light at the growth irradiance (120 μmol m^−2^ s^−1^) and three minutes of darkness. Light was provided by a halogen bulb *via* a liquid light guide. Masses 32, 36, 44 and 45 were monitored with a Sentinel Pro magnetic sector mass spectrometer (Thermo Fisher, USA) allowing the calculation of O_2_ evolution by PSII (mass 32), O_2_ consumption by terminal oxidases and Mehler-type pathways (mass 36), CO_2_ production by mitochondrial activity [minus internal CO_2_ recaptured (mass 44)] and CO_2_ fixation by Rubisco [minus internal CO_2_ recaptured (mass 45)]. Data processing was based on concepts and methods described by [36].

### Transcriptomic meta-analyses

Gene expression data was acquired from ArrayExpress E-MTAB-662 (*rcd1*) [52], E-GEOD36011 (antimycin A) [53] and E-GEOD-9719 (2-hr hypoxia) [54, 55]. Genes that both showed at least a 2-fold change and had a statistical significance of p<0.05 were considered as differentially expressed and were categorised as up- or down-regulated based on the direction of the change. The overlap of multiple gene lists was analysed using Venn diagrams. Pairwise Fischer’s exact test was performed on the gene lists.

### Feeding with ^14^C glucose and analysis of metabolic fluxes

^14^C glucose labelling, fractionation and analysis of metabolic fluxes were performed as described in [9]. Arabidopsis leaf discs were incubated for 150 min in light with 5 mL of 10 mM MES-KOH (pH 6.5) containing 1.85 MBq/mmol [U-^14^C] glucose (Hartmann Analytic) in a final concentration of 2 mM. Leaf discs of the dark experiment were incubated similarly but under the green light. Samples were washed with distilled water, harvested and kept at −80 °C for further analysis. The evolved ^14^CO_2_ was collected in 0.5 mL of 10% (w/v) KOH. Samples were extracted, fractioned and metabolic fluxes were analysed according to [9]. Material from frozen leaf discs was extracted in a two-step ethanolic extraction of 80% (v/v) and 50% (v/v). Supernatants were combined, dried and resuspended in 1 mL of water [56, 57]. The soluble fractions were separated into neutral, anionic, and basic fractions by ion-exchange chromatography as described in [57]. 2.5 mL of the neutral fraction were freeze-dried and resuspended in 100 μL of water for further enzymatic digestions as described in [58]. Phosphate esters of the soluble fractions were measured as in [9] and starch of the insoluble fractions was measured as described in [56]. Calculation of the fluxes was performed according to the assumptions described by Geigenberger et al., [59] and [60].

### Thiol-specific labelling of protein extracts

Thiol-specific labelling of protein extracts was done and interpreted as described in [9].

### Measurement of ATP synthase activity by electrochromic shift

Measurement of ATP synthase activity was done as described in [46].

## Acknowledgements

We thank Dr. Nikolai Belevich for constructing the hypoxia chamber, and Prof. Frank Van Breusegem for providing the *ANAC013* overexpressor line. We are grateful to Dr. Michael Wrzaczek and Dr. Mikael Brosché for critical and helpful comments on the manuscript, to Dr. T. Matthew Robson for suggesting a few important synonyms, and to Dr. Pedro Aphalo for the help in optimizing the anaerobic treatment assay.

## Supplementary figure legends

**Supplementary Figure 1. Gas exchange in wild type and *rcd1* leaf discs as monitored by MIMS.** Wild type is in the left column, *rcd1* is in the right column.

**Supplementary Figure 2. The response of *rcd1* to MV is related to NPQ and is “reset” in darkness.** Chlorophyll fluorescence kinetics observed in *rcd1* in response to MV was suppressed by the proton gradient inhibitor nigericin (A) and in the *rcd1 npq4* double mutant (B), suggesting that the dynamics of Fm’ was related to NPQ. The reads are normalized to Fo. (C) Introducing dark periods (“d”) in the course of light exposure temporarily restored NPQ in MV-treated *rcd1*, indicating that physiological activity of MV in this mutant was reversibly inhibited by light.

**Supplementary Figure 3. The combined effects of MV and hypoxia on PET**. Kinetics of chlorophyll fluorescence during exposure of leaf discs to low light after 15-min flushing of nitrogen gas in darkness. Pre-treatment with MV led to quenched chlorophyll fluorescence in all the tested lines. The presence of this effect in the *npq4*, the *ptox* and the *stn7* mutants suggested that it was not mediated by NPQ, PTOX chloroplast terminal oxidase, and chloroplast state transitions, accordingly. All reads are normalized to Fo obtained under dark-adapted hypoxic conditions.

**Supplementary Figure 4. Response to MV is sensitive to hypoxia in MDS-inducing perturbations other than the *rcd1* mutant**. (A) Pre-treatment of wild-type plants with 2.5 μM antimycin A (AA) makes chlorophyll fluorescence under hypoxia insensitive to MV. This makes AA-treated Col-0 similar to *rcd1*. The reads are normalized to Fo obtained under dark-adapted hypoxic conditions. (B) Quantification of Fs obtained in the experiment shown in (A). The experiment was performed three times with similar results. (C) Similarly to *rcd1*, in *ANAC013* overexpressor line chlorophyll fluorescence under hypoxia was insensitive to MV. The curves are double normalized to fluorescence at Fo and Fi (20 μsec and 40 msec, accordingly). The experiment was performed three times with similar results.

**Supplementary Figure 5. Activity of ATP synthase is unaltered in *rcd1* under standard growth conditions**. To find out whether the activity of the chloroplast ATP synthase was altered in *rcd1* under light-or dark-adapted growth conditions, we performed spectroscopic measurements of thylakoid proton motive force essentially as described in [46]. The decay rate of this parameter after the light flash is proportional to proton conductivity of the ATP synthase. The decay was rapid in light-adapted leaves where ATP synthase was fully activated, whereas dark incubation lead to inactivation of ATP synthase (red and black curves, accordingly). As expected, dark inactivation was less pronounced in the *NTRC* overexpressor line characterized by increased activity of the ATP synthase. In these conditions, the *rcd1* mutant was indistinguishable from the wild type.

