## Supplementary figures for "Increased expression of mitochondrial dysfunction stimulon genes affects chloroplast redox status and photosynthetic electron transfer in Arabidopsis"

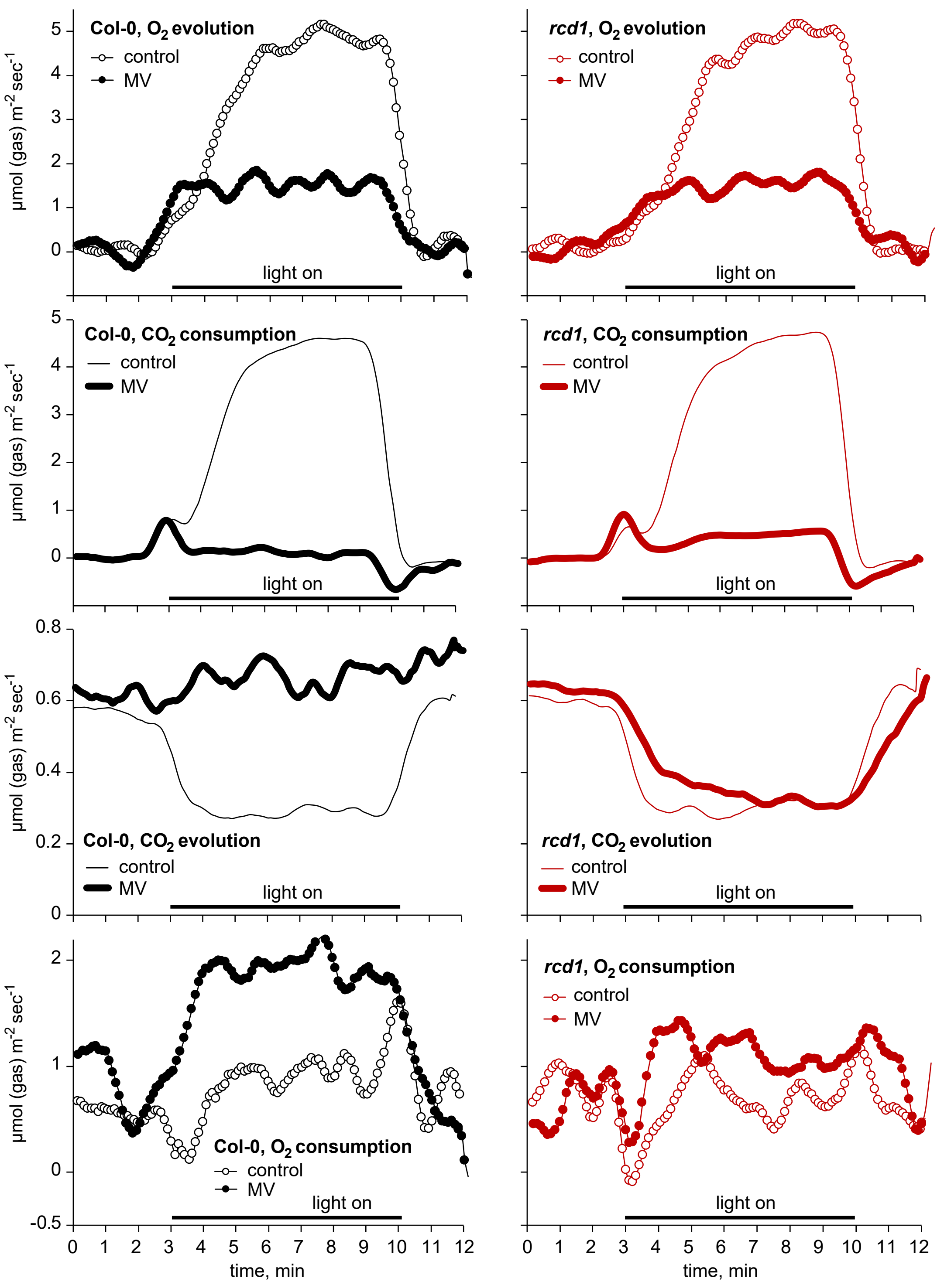


**Supplementary Figure 1. Gas exchange in wild type and *rcd1* leaf discs as monitored by MIMS.** Wild type is in the left column, *rcd1* is in the right column.

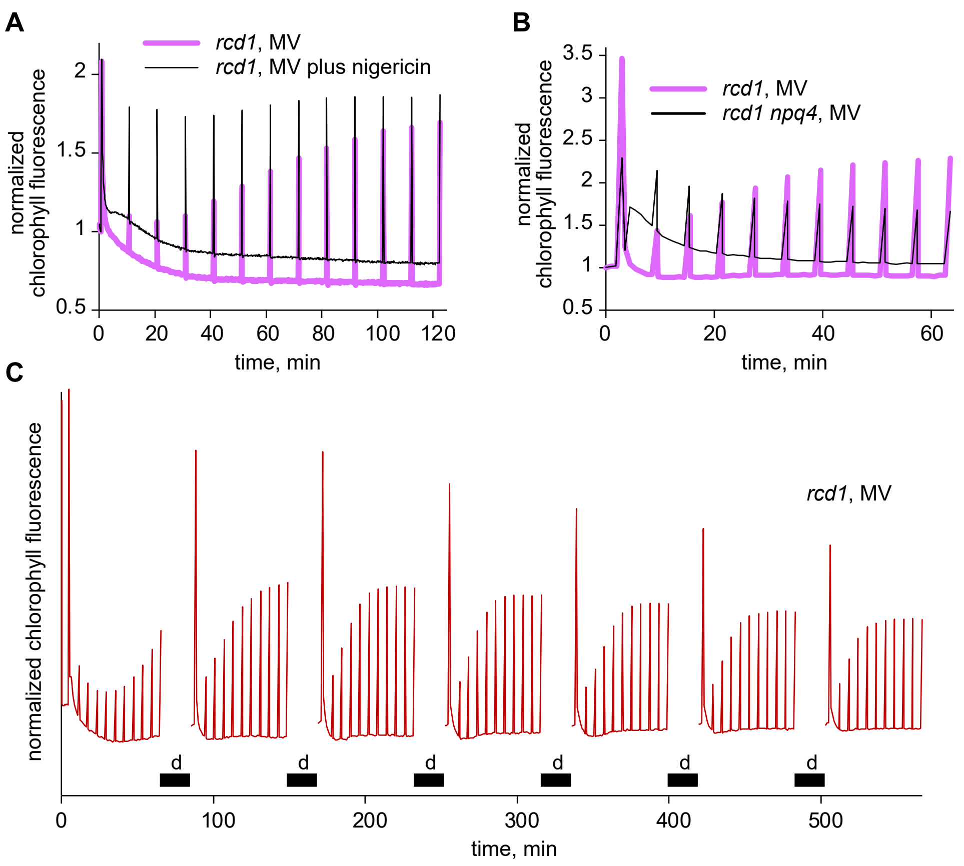


**Supplementary Figure 2. The response of *rcd1* to MV is related to NPQ and is "reset" in darkness.** Chlorophyll fluorescence kinetics observed in *rcd1* in response to MV was suppressed by the proton gradient inhibitor nigericin (A) and in the *rcd1 npq4* double mutant (B), suggesting that the dynamics of Fm' was related to NPQ. The reads are normalized to Fo. (C) Introducing dark periods ("d") in the course of light exposure temporarily restored NPQ in MV-treated *rcd1*, indicating that physiological activity of MV in this mutant was reversibly inhibited by light.

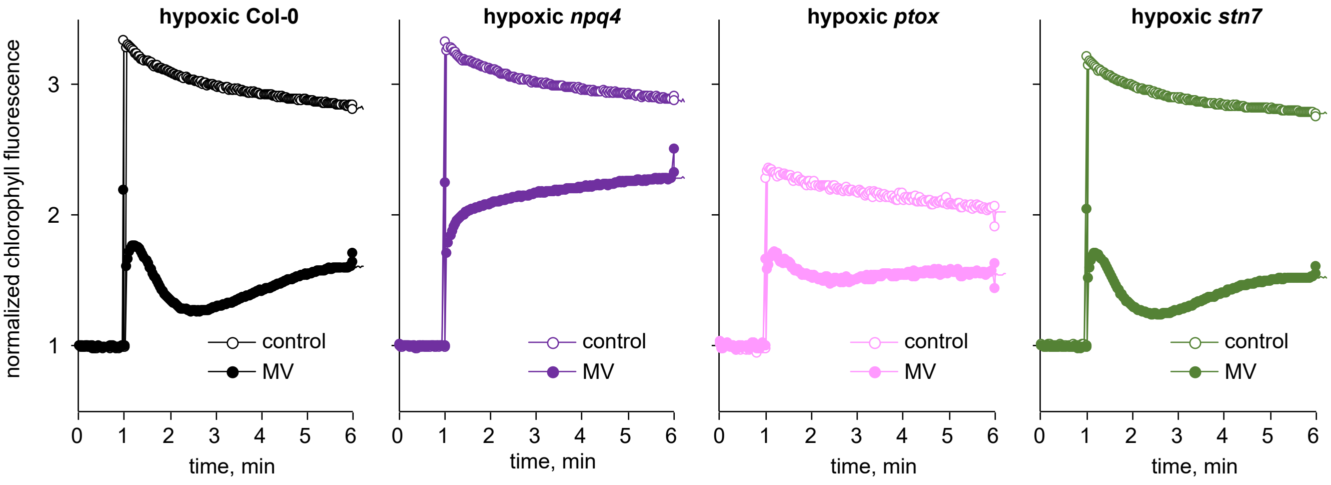


**Supplementary Figure 3. The combined effects of MV and hypoxia on PET.** Kinetics of chlorophyll fluorescence during exposure of leaf discs to low light after 15-min flushing of nitrogen gas in darkness. Pre-treatment with MV led to quenched chlorophyll fluorescence in all the tested lines. The presence of this effect in the *npq4*, the *ptox* and the *stn7* mutants suggested that it was not mediated by NPQ, PTOX chloroplast terminal oxidase, and chloroplast state transitions, accordingly. All reads are normalized to Fo obtained under dark-adapted hypoxic conditions.

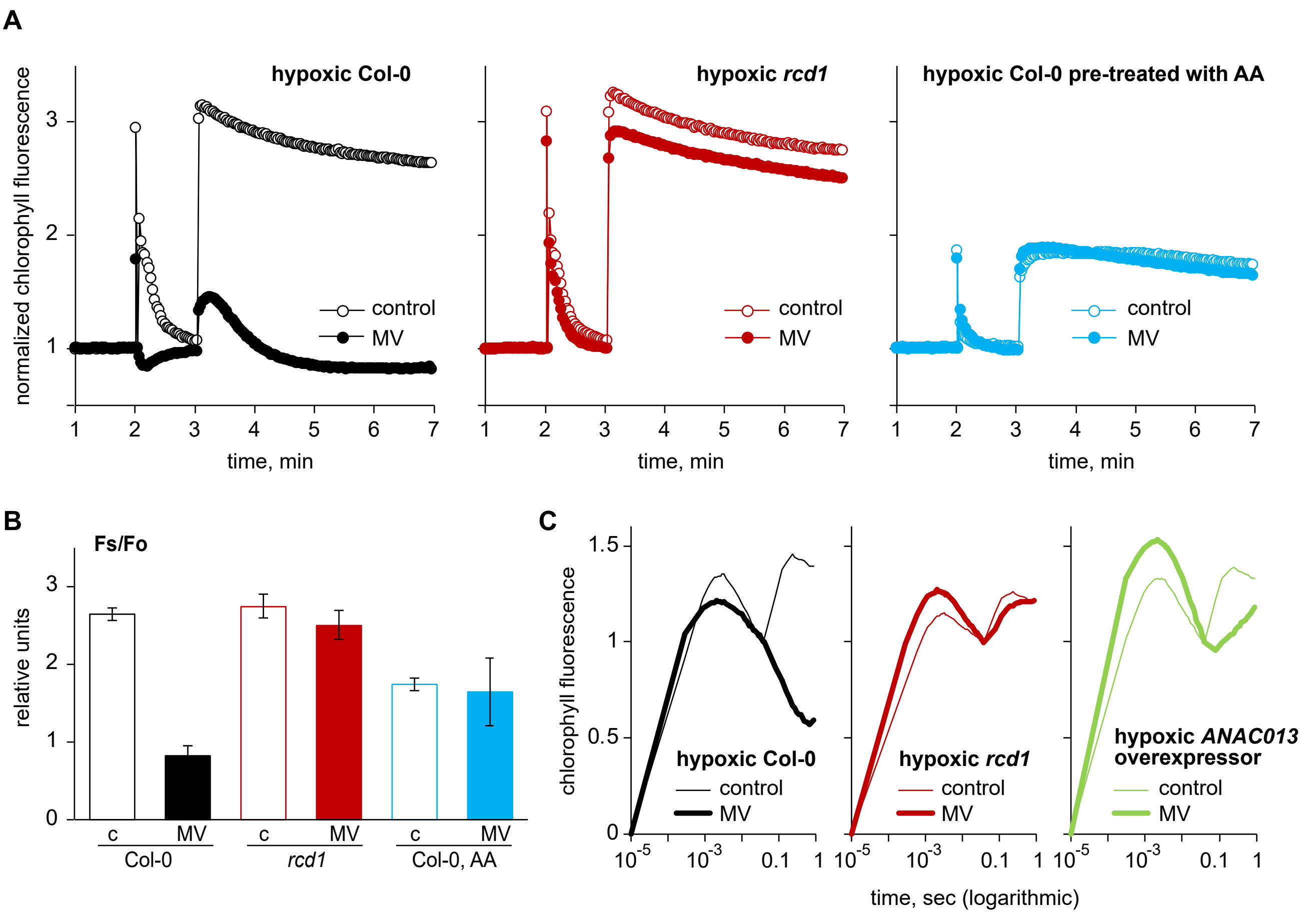


**Supplementary Figure 4. Response to MV is sensitive to hypoxia in MDS-inducing perturbations other than the *rcd1* mutant.** (A) Pre-treatment of wild-type plants with 2.5 µM antimycin A (AA) makes chlorophyll fluorescence under hypoxia insensitive to MV. This makes AA-treated Col-0 similar to *rcd1*. The reads are normalized to Fo obtained under dark-adapted hypoxic conditions. (B) Quantification of Fs obtained in the experiment shown in (A). The experiment was performed three times with similar results. (C) Similarly to *rcd1*, in *ANAC013* overexpressor line chlorophyll fluorescence under hypoxia was insensitive to MV. The curves are double normalized to fluorescence at Fo and Fi (20 µsec and 40 msec, accordingly). The experiment was performed three times with similar results.

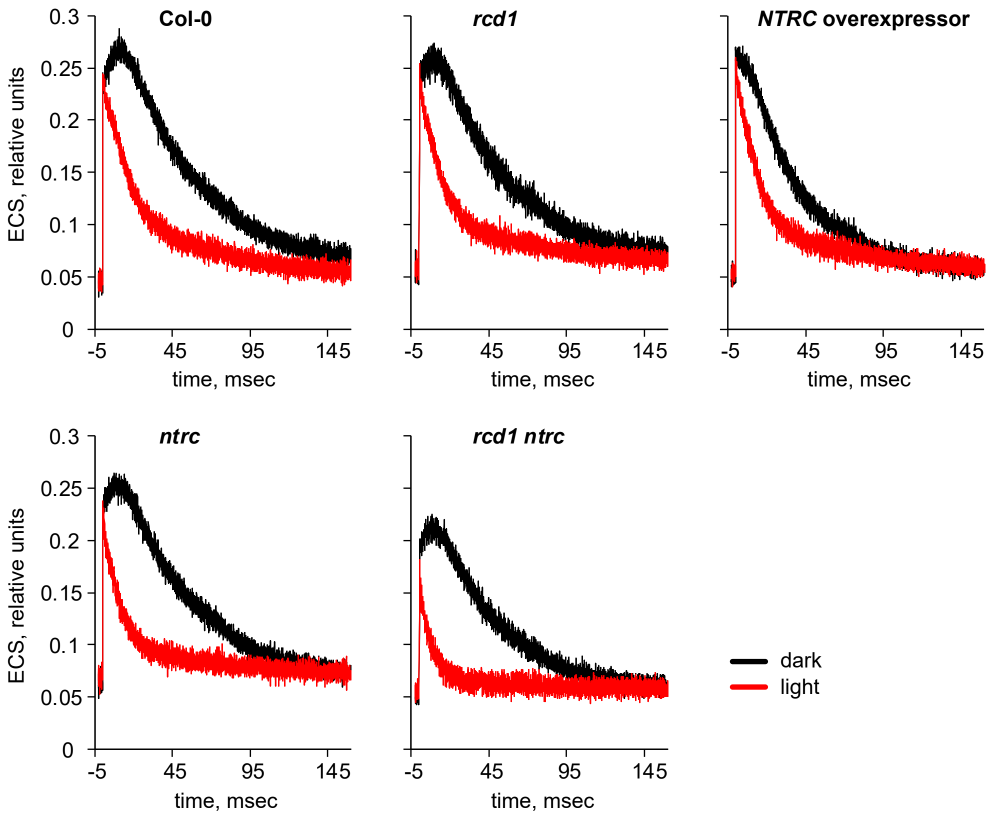


**Supplementary Figure 5. Activity of ATP synthase is unaltered in *rcd1* under standard growth conditions.** To find out whether the activity of the chloroplast ATP synthase was altered in *rcd1* under light-or dark-adapted growth conditions, we performed spectroscopic measurements of thylakoid proton motive force essentially as described in [46]. The decay rate of this parameter after the light flash is proportional to proton conductivity of the ATP synthase. The decay was rapid in light-adapted leaves where ATP synthase was fully activated, whereas dark incubation lead to inactivation of ATP synthase (red and black curves, accordingly). As expected, dark inactivation was less pronounced in the *NTRC* overexpressor line characterized by increased activity of the ATP synthase. In these conditions, the *rcd1* mutant was indistinguishable from the wild type.
